# Oligomerization of the *Clostridioides difficile* Transferase B Component Proceeds through a Stepwise Mechanism

**DOI:** 10.1101/2025.05.06.652354

**Authors:** Robin M. Mullard, Michael J. Sheedlo

## Abstract

*Clostridioides difficile* is a gram-positive, pathogenic bacterium and is currently the leading cause of hospital-acquired, infectious diarrhea in the United States. During infection, *C. difficile* produces and secretes up to three toxins called Toxin A, Toxin B, and the *C. difficile* transferase (CDT). While Toxin A and Toxin B are thought to drive the pathology associated with the disease, strains that produce CDT have been linked to increased disease severity, higher rates of infection recurrence, and increased incidence of mortality. A basic understanding of how CDT intoxicates host cells has emerged over the past two decades and includes a framework that relies on the oligomerization of the components that comprise CDT to promote cellular intoxication. Although several key steps of this process have been biochemically described, a clear, molecular description of toxin assembly has not been resolved. We have collected cryogenic electron microscopy (Cryo-EM) data of purified, recombinant CDT. From these data, we have generated several structural snapshots of the toxin, including a series of structures that correspond to intermediates that form during oligomerization. These structures provide insight into the mechanism underlying toxin assembly and highlight a role for structural plasticity during this process. We have also shown that these partially assembled toxins are equally potent in cytotoxicity assays supporting this model in a cellular context. Finally, we show that CDTb oligomers are stabilized by CDTa and assembly is triggered by hydrophobic molecules.

**Author Summary:** *Clostridioides difficile* is the leading cause of hospital-acquired, infectious diarrhea in the United States. Strains of *C. difficile* that produce a toxin known as the *C. difficile* Transferase (CDT) are linked to more severe presentations of the disease in clinical settings. Despite its apparent importance, a molecular model of CDT intoxication remains undeveloped. Reported here are several Cryo-EM structures that describe the mode of CDT assembly which occurs through the stochastic and sequential addition of proteins that comprise the toxin.

## Introduction

*Clostridioides difficile* is a pathogenic, gram-positive bacterium and is the leading cause of hospital-acquired, infectious diarrhea in the United States (1). The pathology that accompanies *C. difficile* infection is induced by the activity of toxins produced by the bacterium that target the large intestine known as Toxin A (TcdA), Toxin B (TcdB), and the *C. difficile* transferase (CDT) (2). The number and type of toxins produced by *C. difficile* varies from one strain to the next, with some pathogenic strains producing only Toxin B (3). However, the strains observed most frequently in clinical settings produce all three toxins and are linked to more severe symptoms, modulation of the host immune response, higher rates of disease recurrence, and increased incidence of mortality (4–7). This observation has led to the suggestion that all three toxins act synergistically to promote severe disease and a renewed interest in understanding how all three toxins contribute to these phenotypes. Of the three toxins, CDT is the least studied and progress in understanding its contribution to disease has been hindered by an incomplete molecular model of intoxication.

CDT is a member of the Iota family of protein toxins which also consists of two homologous toxins produced by *Clostridium perfringens* (the Iota toxin) and *Clostridium spiroforme* (*C. spiroforme* Transferase, CST) (8–11). All three Iota toxin family members are assembled from two polypeptide chains termed the “A” subunit (CDTa, Ia, and CSTa) and “B”, subunit (CDTb, Ib, and CSTb). The A subunit of all three toxins encodes three features 1) an N-terminal unstructured region, 2) a pseudo-ADP-ribosyltransferase domain (pADPRT) and 3) an ADP-ribosyltransferase domain (ADPRT) (Fig S1A) (12,13). All three B subunits are presumed to contain five domains that are involved in regulation (D1), oligomerization (D2 and D3), and host recognition (D3’ and D4) (Fig S1B) (14–16). Often discussed in concert with the Iota toxins are the C2 toxin of *Clostridium botulinum* and the anthrax toxin produced by *Bacillus anthracis*, though it is important to note key differences exist in the sequences and structures of C2 and anthrax toxin that suggest they are mechanistically distinct from the Iota toxins (11,14–17).

The accepted model of CDT intoxication has been derived from studies conducted on all three Iota toxins and supplemented with observations made on the C2 and anthrax toxins due to their perceived similarities. The mechanism of intoxication starts with the secretion of CDTa and CDTb from the bacterium at the site of infection (10). CDTb is then thought to localize to host cells via interactions with a host cell receptor known as Angulin-1 (via the D4 domain), and possibly glycans (via the D3’ domain) where it is activated by proteases produced by the host cell (14,18). Once proteolyzed, the mature form of CDTb forms a large heptameric structure referred to as the prepore which binds one molecule of CDTa to generate the toxic moiety, CDT (14,16,19–21). CDT enters host cells by receptor-mediated endocytosis where the environment within the maturing endosome triggers a structural change within CDTb that leads to a repositioning of the D2 domain and the formation of a membrane-spanning pore (21). The channel formed by CDTb is then thought to serve as a conduit through which CDTa enters the host cell cytoplasm. Inside the host cell, CDTa modifies actin and the actin related protein 2/3 leading to cytoskeletal deformation and the protrusion of microtubules that interact with the bacterium at the site of infection (Fig S1C) (10,22).

Although the formation of oligomeric structures is a critical step in the intoxication pathway of CDT, little is known about how the toxic moiety assembles (21). Instead, anthrax toxin has been used as a surrogate for understanding CDT assembly as it is the closest relative for which experimental data detailing the mechanism of assembly have been proposed. In the case of anthrax toxin, the delivery component (a protein known as the protective antigen) has been shown to form dimeric structures upon interaction with the host cell receptor, ANTXR (23). These dimers are presumed to associate to form larger structures culminating in the formation of oligomers containing either seven (heptameric) or eight (octameric) copies of the protective antigen (24). Unlike CDT, anthrax toxin delivers two enzymes into the cytoplasm of host cells known as the edema and lethal factors (25). The step at which the edema and lethal factors associate with the protective antigen to form the anthrax toxin is not yet known. However, structural studies have uncovered similar binding modes for the two components that rely on an interface formed across two adjacent protective antigen protomers suggesting the immature dimer may recognize these cargo during toxin maturation (26,27). It is not currently clear if, or how, the assembly pathway of anthrax toxin might translate to CDT. Indeed, significant differences in the structures and sequences of the enzymatic cargo of the two toxins result in differential arrangements of the A and B components in the assembled toxins (Fig S1D) (15,19,20). Similarly, the receptor binding domains of CDTb and the protective antigen adopt radically different configurations, suggesting CDT may require a different trigger to stimulate oligomerization (Fig S1E) (14,16,17). The goal of this study was to elucidate the mechanism of CDT assembly. To that end, we have generated cryogenic electron microscopy (Cryo-EM) structures of several forms of CDTb and propose the toxin assembles through the simple addition of CDTb protomers. Moreover, our data indicate that assembly intermediates can form interactions with CDTa, suggesting the enzymatic component spontaneously associates with the intermediates that form during assembly. We also note that interactions with CDTa may drive CDTb oligomerization as the configuration of the CDTa-bound oligomer more closely resembles that of the assembled toxin. Finally, we provide evidence that our proposed model occurs in a cellular context and illustrate the role of hydrophobic molecules in driving toxin assembly.

## Results

### Cryo-EM Analysis of CDTb Oligomers Suggests a Stepwise Mechanism of Assembly

To probe the assembly mechanism of CDT, CDTb was recombinantly purified and proteolytically activated with trypsin which led to the spontaneous formation of oligomeric structures as previously described (13–16,19,20). CDTa was then mixed with CDTb at a molar ratio of 1:1 (CDTa:CDTb) and the sample vitrified for high resolution data collection via Cryo-EM (Fig S2A). Subsequent analysis led to the identification of monomeric CDTb, several CDTb oligomeric intermediates (including dimers, trimers, tetramers, pentamers, hexamers, and heptamers), and large oligomeric structures (including heptamers and octamers) in this sample (Fig 1A-B, Fig S2B). Notably, the oligomeric intermediates displayed a pattern of association wherein the protomers were arranged about a central pseudosymmetry axis running through the center of the oligomer, consistent with that of the symmetry axis observed in the mature toxin. The number of particles representing each class were enumerated during subsequent classification and yielded significantly different particles counts for the oligomeric intermediates (60,753 monomeric, 330,267 dimeric, 76,576 trimeric, 43,725 tetrameric, 14,329 pentameric, 2,688 hexameric, and 1,456 heptameric CDTb assemblies were observed, Fig S2C). We note that the number of particles larger than the monomeric structure can be fit to an exponential decay function when plotted against the number of protomers contained within each structure (R^2^ = 0.9943) indicating the assembly of CDTb oligomers is a stepwise process (Fig 1C). Similarly, the number of symmetric particles in this dataset were not equal, with the heptameric structure ∼44-fold more abundant than the octameric form (Fig 1D).

**Fig 1.**
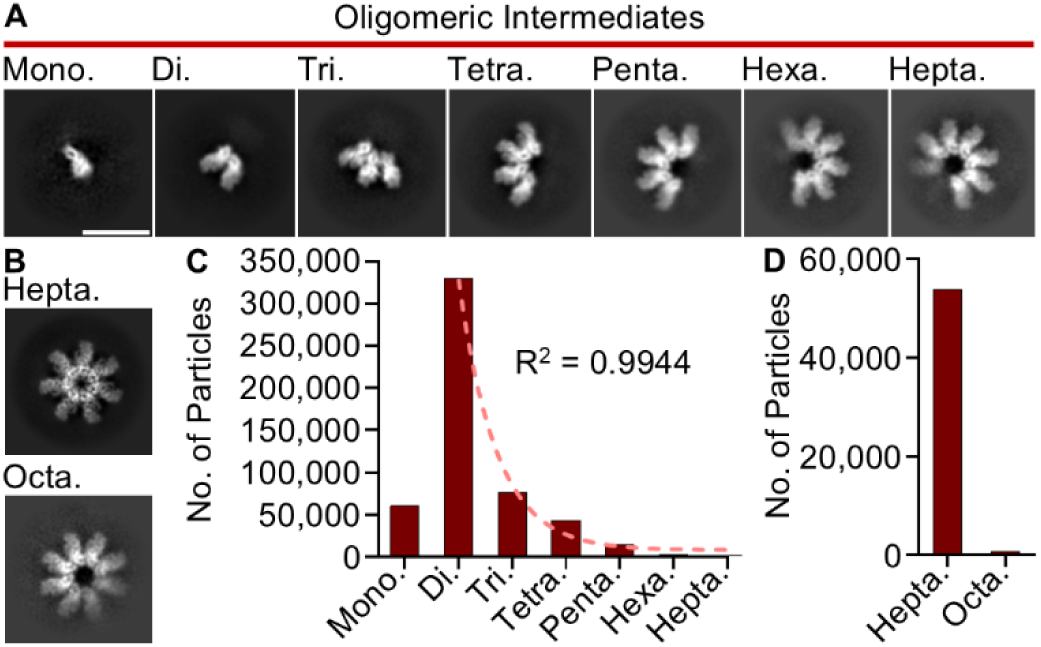
Oligomeric CDTb Structures Identified by Cryo-EM. (**A**) Two-dimensional classes of seven CDTb structures corresponding to oligomeric intermediates. Mono. – monomer, Di. – dimer, Tri. – trimer, Tetra. – tetramer, Penta. – pentamer, Hexa. – hexamer, and Hepta. – heptamer. Scale bar represents 100 Å and is the same across all seven classes. (**B**) CDTb symmetric structures corresponding to heptamers (Hepta.) and octamers (Octa.) as observed in the Cryo-EM dataset. (**C**) The number of particles identified that correspond to each oligomeric intermediate. Particle counts for structures larger than a monomer were fit to an exponential decay function. (**D**) The number of particles classified as either a symmetric heptamer or octamer.

Three-dimensional maps were reconstructed for all oligomeric structures observed within this dataset, except the asymmetric heptamer which lacked enough particles to generate a reliable reconstruction. However, due to significantly lower particle counts for the larger oligomeric intermediates, a decrease in resolution was observed for all intermediate structures containing more than three protomers. Still, maps of modest global resolution could be reconstructed for the CDTb dimer (3.74 Å), trimer (4.08 Å), tetramer (4.08 Å), and symmetric heptamer (3.27 Å) which enabled map refinement and model construction (Fig S2D). Further classification and refinement of the CDTb dimer led to a map with a final resolution of 3.56 Å, with local resolutions consistent throughout much of the map (Fig S3A, B). Modeling of the CDTb dimer resulted in the assignment of two chains that we call ‘Chain A’ and ‘Chain B’, both of which were observed in a prepore-like configuration with three additional nodes of density that we modeled as calcium ions in agreement with previous studies (Fig 2A, Fig S3C, D). The modeled portions of the two chains are nearly identical in structure with a low backbone RMSD (0.945 Å), though differences in the residues that could be modeled do exist (Fig S3E, Table S1). The receptor binding domain (D4) was not observed in either chain in contrast to previous structures though additional density is present, indicating the D4 domain remains flexible during toxin assembly (Fig S3F). Chains A and B are arranged in positions and orientations that are similar to that observed in the prepore structure, though Chain B is rotated ∼40° about the central pseudosymmetric axis in the dimeric intermediate, as opposed to ∼51° in the structure of the prepore, illustrating structural plasticity at the dimeric interface during assembly (Fig 2A). The surface formed between Chains A and B accounts for ∼1,200 Å^2^ of buried surface area and relies primarily on structures formed by residues 411-423, 477-484, 504-513, and 534-541 from Chain A and residues 237-240, 278-290, 427-434, 439-451, and 490-498 from Chain B for assembly (Fig S3G). All residues contained within this interface are also observed in the structure of the symmetric heptamer, reconstructed from the same dataset. One site of interaction comprised of structures formed by residues 384-388 and 456-463 of Chain A and residues 314-318 and 451-454 of Chain B was observed in the previously defined structure of the CDTb prepore but is not found to interface in the oligomeric intermediate suggesting it is only engaged in the fully formed toxin (Fig 2B).

**Fig 2.**
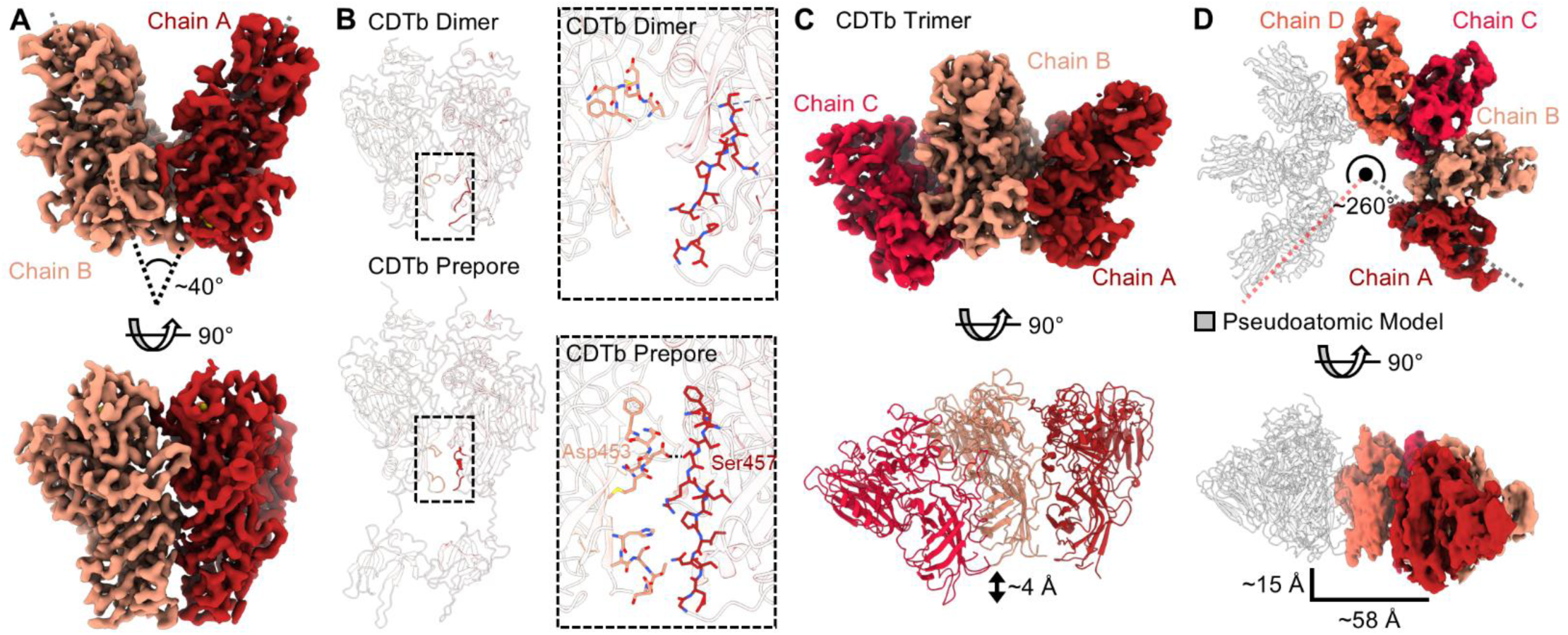
Structures of the CDTb Oligomeric Intermediates. (**A**) A map of the CDTb dimer was reconstructed to a resolution of 3.56 Å and consists of two chains termed Chain A (scarlet) and Chain B (salmon). The angle describing the relative orientations of Chains A and B with respect to the central symmetry axis is indicated (**B**) The interface facilitating the formation of the CDTb dimer (top) compared to the dimeric interface observed in the CDTb prepore (PDB 6O2N, bottom). A magnified view of each interface is shown to the right of each image. (**C**) A map of the CDTb trimer was reconstructed to 4.19 Å resolution (top) and consists of three chains defined as Chain A (scarlet), Chain B (salmon), and Chain C (red). A lateral displacement of Chain C by ∼4 Å was observed with respect to Chains A and B. (**D**) A model of the CDTb asymmetric heptamer constructed from the 7.01 Å map of the CDTb tetramer (top). This model illustrates a lateral displacement of protomeric units that, when propagated, results in a shift of ∼15 Å shift in the lateral dimension and a ∼58 Å gap between terminal protomers resulting in a ‘lock-washer’ shape.

A map of the CDTb trimer was refined to a final resolution of 4.19 Å. Although the nominal resolution of the final map is lower than the initial reconstruction, the final map does not suffer from problems arising from preferred orientation as is apparent in the initial map (Fig S4A). The model of the dimer was used as a reference to generate a molecular model of the trimeric intermediate (Fig S4B, Table S1). We note a similar arrangement of Chains A and B within this structure and the presence of a third chain, ‘Chain C’, positioned next to Chain B, opposite of Chain A (Fig 2C). As observed in the dimeric structure, the orientations of all three chains are rotated compared to the assembled, symmetric heptamer with an observed rotation of ∼40° between Chains A and B and ∼43° between Chains B and C (Fig S4C). The interface between Chains A and B and Chains B and C mirror those of the dimeric intermediate and do not indicate the presence of additional recognition elements (Fig S4D). Interestingly, a helical pitch is also observed within the trimer wherein Chain C is positioned ∼4 Å out of plane when compared to the symmetric structure (Fig 2C). A map of the tetrameric intermediate was refined after rebalancing particles to limit issues arising from preferred orientation leading to a map of 7.01 Å resolution. The helical nature of the oligomeric intermediates is also observed in the model of the tetrameric intermediate which illustrates a separation of ∼46 Å across the tetramer (Fig S4E). When propagated about the asymmetric heptameric structure, this would account for a total separation of ∼15 Å in the vertical dimension and ∼58 Å in the horizontal direction forming a ‘lock-washer’ like structure and indicating the structural plasticity observed across the oligomerization interface is not restricted to two-dimensional space (Fig 2D).

### CDTa Binding to CDTb Oligomeric Intermediates Indicates Sporadic Assembly of CDT

During classification, we noted a population of the CDTb dimeric assembly intermediate bound to CDTa. Despite a significant number of particles observed in this complex, the three-dimensional reconstruction was limited to modest resolution (4.57 Å) due to issues arising from preferred orientation (Fig S5A). Nonetheless, we have successfully assembled a pseudoatomic model of this interaction (Fig 3A, Fig S5B). The interface formed between CDTa and the CDTb dimer is facilitated by interactions with both chains of CDTb (buried surface areas of ∼241 Å^2^ and ∼122 Å^2^ were observed for Chains A and B, respectively) (Fig S5C). Though we note the resolution of the map hindered the assignment of residues at the interface, the CDTb dimer is confirmed to adopt a prepore-like configuration as observed in the apo structure (RMSD of 0.657 Å) and interacts with CDTa through a binding mode that is similar to that previously described (RMSD of 2.203 Å, Fig S5D).(13,19,20) We also uncovered a subpopulation of the CDTb trimer bound to CDTa but note that these particles are present in a significantly lower abundance, indicating the CDTa bound dimer is preferred (Fig 3B, Fig S5E). Still, we were able to generate a low-resolution map (8.15 Å) and assemble a model of the CDTb trimer bound to CDTa. The resulting structure illustrates rotations of Chains B and C of ∼4° and ∼9° when compared to the apo structure, respectively, that allows for interactions with CDTa in both chains (Fig S5F). This results in an orientation of Chain C that closely resembles CDTb protomers as they are arranged in the symmetric heptamer (Fig 3C). Interestingly, no classes pertaining to the CDTa-bound tetrameric, pentameric, hexameric, or heptameric oligomeric intermediates were observed in this dataset.

**Fig 3.**
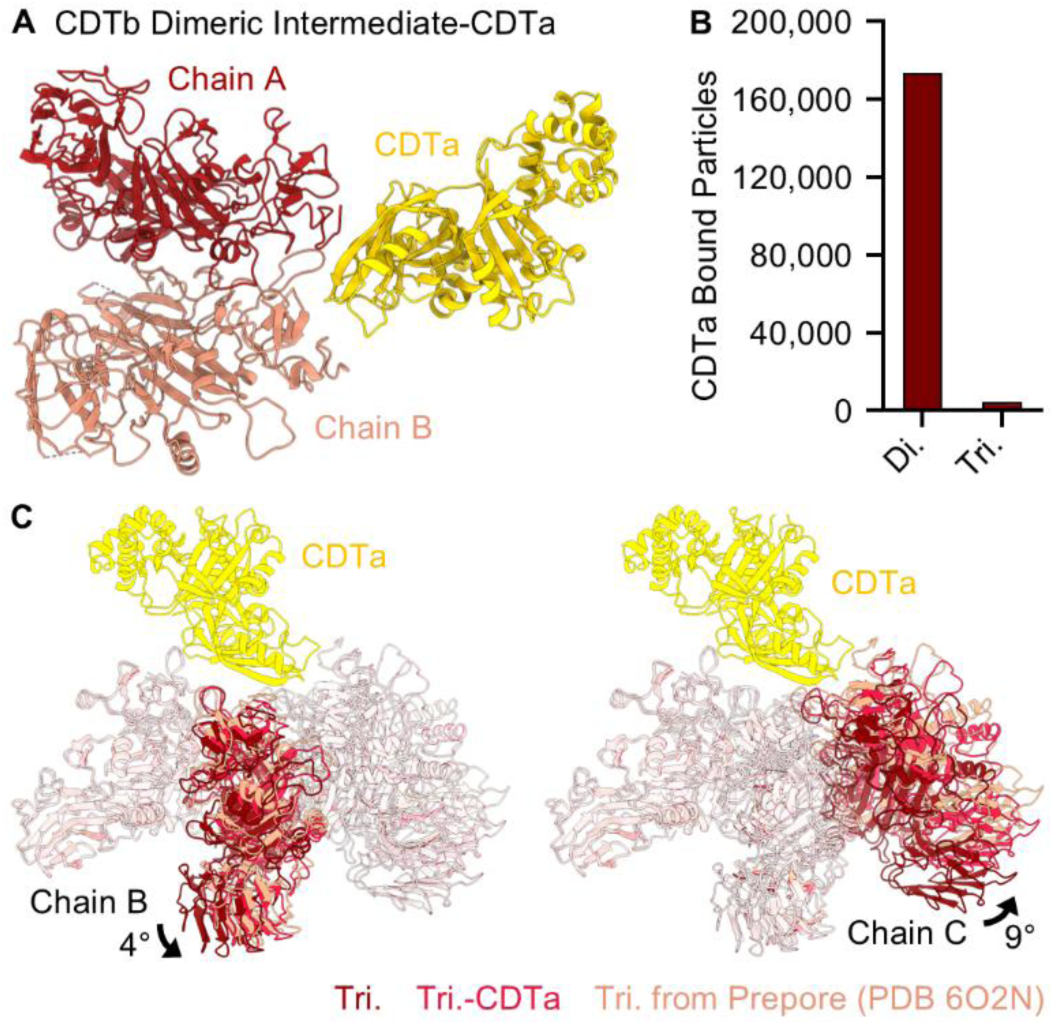
Structure of CDTb Assembly Intermediates Bound to CDTa. (**A**) A structure of the CDTb dimeric assembly intermediate was constructed that includes two chains of CDTb termed Chain A (scarlet) and Chain B (salmon) and one chain of CDTa (gold). (**B**) The relative number of CDTa bound dimeric and trimeric assembly intermediates observed in this dataset. (**C**) The relative orientations of Chains B and C within the trimeric intermediate shift to contact CDTa, resulting in a structure that resembles that of the assembled, symmetric prepore.

### CDTb Assembly Intermediates Display Equal Potency in a Cellular Model

We next developed a protocol to isolate CDTb oligomeric intermediates to test their potency in a cellular intoxication model. The oligomeric building blocks were purified via size exclusion chromatography using which we isolated five fractions (denoted by number, 4-8) of differing molecular weights (Fig 4A). The presence of CDTb oligomers were confirmed for all fractions via SDS-PAGE as the oligomeric structures formed during intoxication are resistant to denaturation by SDS (Fig S6A). All five of these fractions were mixed with excess CDTa, applied to Caco-2 cells in tissue culture, and the development of round cells was used to quantify cellular intoxication as we have previously described (13). We note that, under these conditions, all fractions were found to be active, indicating that these oligomeric building blocks are functional. To assess the potency of each isolate, we titrated all five fractions on Caco-2 cells ranging in protomer concentration from 7,000 pM to 56 pM. All fractions except for fraction four (which is presumed to also contain unfolded and aggregated particles), were found to be equally potent on Caco-2 cells after a six-hour incubation (Fig 3B, Fig S6B). Because we anticipate larger oligomeric building blocks would bind to multiple receptors, we expected the larger fragments would have a higher affinity for the cell surface and thus exhibit higher potencies. Because this phenomenon was not illustrated in our cell rounding assays, we reasoned that the quantity of each oligomeric intermediate formed in solution is dependent on the concentration of protomeric CDTb in the sample. To demonstrate this phenomenon, we isolated fractions five and eight which were resubjected to size exclusion chromatography. Upon separation, we note the appearance of larger oligomeric intermediates in fraction eight indicating the oligomeric intermediates exist in equilibrium (Fig S7A). Conversely, the particles isolated in fraction five retained a high molecular weight, suggesting the symmetric heptamer does not spontaneously disassemble upon dilution (Fig S7B).

**Fig 4.**
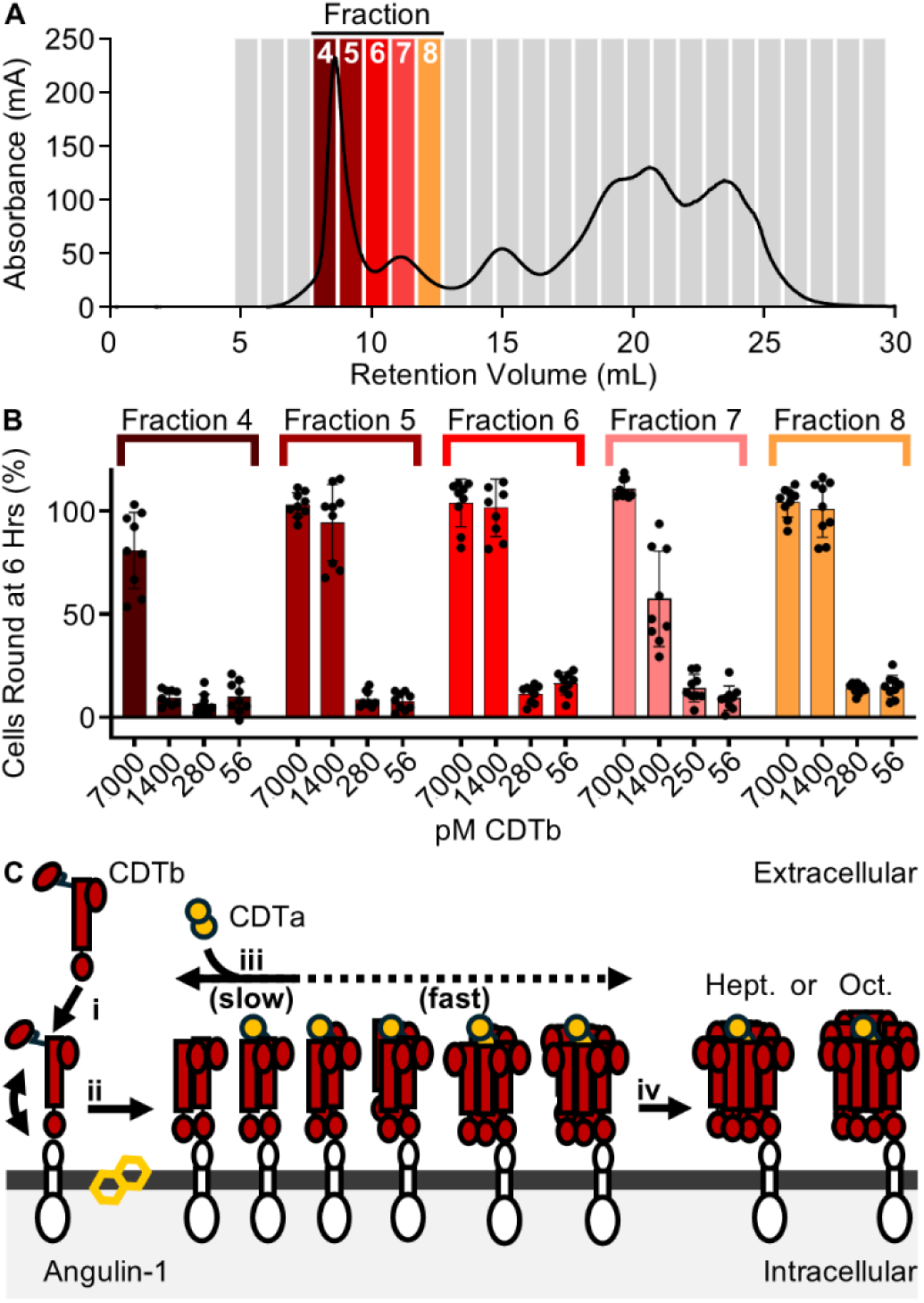
Intoxication by CDTb Oligomeric Intermediates. (**A**) CDTb oligomeric intermediates were fractionated via size exclusion chromatography to yield five distinct samples termed fractions 4-8. (**B**) The intoxication efficiency of the fractions containing CDTb oligomeric intermediates on Caco-2 cells in tissue culture after a six-hour incubation. Error bars represent standard deviation. (**C**) A model of CDTb toxin formation that illustrates the stepwise assembly of CDTb prepore moieties stimulated by interactions with hydrophobic molecules found in the host cell membrane and potentially CDTa, resulting in the formation of symmetric heptameric and octameric structures.

### CDTa Enhances the Formation of Stable CDTb Heptamers

Because CDTa was found to impact the arrangement of the CDTb trimeric intermediate, we next sought to determine how the inclusion of CDTa impacts the distribution of oligomeric intermediates in solution. Using size exclusion chromatography, we analyzed samples containing CDTa, CDTb oligomeric intermediates, and CDTb oligomeric intermediates in the presence of CDTa. We note that the sample containing both CDTa and the CDTb oligomeric intermediates yielded a small, but notable, high molecular weight peak at a position corresponding to the heptameric structure that is not present in the samples in isolation (Fig S7C). Intriguingly, the concentration of the large molecular weight species remains low, suggesting CDTa does not significantly accelerate oligomerization but may, instead, play a role in locking CDTb in the heptameric configuration.

### CDTb Oligomerization is Stimulated by Hydrophobic Molecules and CDTa

To define the driver of CDTb oligomerization, we used an SDS-PAGE based assay to track oligomer formation in solution. Using this assay, we first confirmed that CDTb oligomerization is slow in solution with no oligomer formed over a period of 60 minutes (Fig S8A). Following previous reports that ethanol stimulates oligomerization in vitro, we demonstrated that CDTb oligomerization is dependent upon the concentration of ethanol in solution with a ∼7.5-fold enhanced efficacy of oligomerization observed following the inclusion of 2.5 M ethanol (15,20). We then extended this analysis to a small panel of solvents, including acetone, methanol, ethanol, isopropanol, and dimethyl sulfoxide to define the property of the solvent responsible for this phenomenon. Of the solvents tested, isopropanol was found to be the most effective followed by acetone and ethanol. To assess the possibility that these solvents induce molecular crowding to induce oligomerization we also included polyethylene glycol 8,000 in this assay but note no obvious effect when it is included under these conditions (Fig S8B). We next reasoned that solvent hydrophobicity may drive activity. Indeed, when plotting the hydrophobicity of each solvent against its capacity to induce oligomerization, we note the resulting plot can be fit to an exponential decay function (R^2^ = 0.9374), indicating hydrophobicity drives this phenomenon (Fig S8C). While these solvents are not likely to be present at the site of infection in appreciable concentrations we argue they mimic hydrophobic molecules associated with the host cell membrane which are expected to significantly stimulate CDTb oligomerization in vivo.

## Discussion

Despite its apparent importance in driving the severe symptoms associated with *C. difficile* infection in some strains, a complete description of CDT intoxication has yet to emerge. To fill critical gaps in this pathway, we have leveraged the power of Cryo-EM and present ten unique structures of CDT, eight of which we suggest are correlated to intermediates that are formed during toxin assembly. The data we present here illustrate an assembly mechanism that is dependent upon the sequential association of CDTb protomers to form large oligomeric structures, generally referred to as a prepore (Fig 4C) (14). This mechanism of assembly is markedly different from that proposed for the related anthrax toxin of *B. anthracis* wherein the protective antigen forms dimers upon binding the host cell receptor, ANTXR (23). These dimers are then expected to associate to form larger oligomeric structures culminating in the formation of symmetric structures comprised of either seven (heptamers) or eight (octamers) protomers (23,24). Although the unique pathway we describe for CDT also results in the formation of heptameric and octameric structures, the octameric structures are present at a significantly lower abundance in vitro (∼1.5% and ∼30% in CDT and anthrax toxin, respectively) (24). We expect that this difference is a product of the assembly pathways of the toxins wherein the protective antigen dimers are formed early in the pathway through associations with ANTXR and lead to a higher abundance of octamers formed through the association of four protective antigen dimers. Conversely, CDTb assembly does not appear to be driven by receptor association as the D4 domain was found to remain flexible in solution limiting the effect this interaction may have on toxin assembly. Because the octameric particles are more stable than their heptameric counterparts, it is likely that the assembly mechanism of anthrax toxin has evolved to incorporate the formation of dimers to enhance the formation of the octameric structure in response to the harsh conditions found at the site of infection (22).

Two of the assembly intermediates described in the present work were defined at high resolution (the CDTb dimer and trimer) and allowed us to define the interface that mediates protomer association during toxin assembly. While the surface we describe here is reminiscent of the interface that facilitates interactions between protomers in the assembled, symmetric prepore we note it is not identical. Indeed, residues 456-463 of CDTb remain unengaged and dynamic in the structures we describe here but interface in the symmetric heptamer. Residues 456-463 of CDTb are positioned near alpha helix 5 (α5, formed by residues 464-471) in the three-dimensional structure which was recently found to play a role in the conversion of the CDTb prepore into a membrane-spanning channel or ‘pore’ (28). Intriguingly, the authors present a stepwise mechanism for this conversion as well. It is thus possible that these residues are the last to engage at the interface between adjacent protomers and act as a sensor to trigger pore formation only when the symmetric structure has been effectively assembled. This model would account for previous observations that describe the apparently uncontrolled transition from the prepore state to the pore state as observed throughout the family of Iota toxins (14–16,19,20). It is also plausible that this interaction functions as a lock, preventing the prepore from disassembling once the structure has formed. Our observation that small oligomeric intermediates, but not the assembled symmetric structures, spontaneously assemble and disassemble would support this hypothesis.

Finally, we have elaborated upon the kinetics of prepore assembly and note the apparently slow rate of association. Previous studies have yielded an important clue to the external factors stimulating toxin assembly and identified ethanol as a potent inducer of oligomer formation (20,28). Expanding upon this observation, we show that other solvents can drive this process and define the hydrophobicity of the solvent as the key determinant in its effectiveness. This observation suggests the possibility that hydrophobic molecules present on or within the cellular membrane may precipitate this process in a cellular context. Similarly, the reorientation of CDTb protomers in the assembly intermediates bound to CDTa suggests that CDTa may also induce toxin assembly through the stabilization of the heptameric form of CDTb. The model we present allows for the acceleration of CDTb oligomerization via hydrophobic molecules present in the cellular membrane and the ‘locking’ of CDTb oligomers into the heptameric structure via the inclusion of CDTa (Fig 4C). Future studies into the intoxication pathway should shed light into the mechanism that governs these functions.

## Materials and Methods

### Purification of CDTa and CDTb

CDTa_50-463_ (pSLMn0003) and CDTb_45-876_ (pSLMn0014) were transformed into *Escherichia coli* strains denoted BL21 Gold and BL21 RIL, respectively. The recombinant protein was then purified following a protocol that was virtually identical to that previously described (13). CDTb was activated via trypsin cleavage to induce oligomerization before further purification by size exclusion chromatography, separating the sample into 1 mL fractions for further analysis.

### CDTb Assembly Intermediate Intoxication Assay

Human male colorectal carcinoma cells (Caco-2, ATCC HTB-37) were grown in Eagle’s Minimum Essential Media (EMEM) supplemented with 20% fetal bovine serum and maintained in an atmosphere of 5% CO_2_ at 37 ℃. Cells were then plated and grown to 90% confluency. Before conducting intoxication assays, the media was removed and replaced with EMEM supplemented with 20% FBS and Hoechst stain (at a 1:100 dilution) for one hour. After incubation, the stain was removed and the cells washed with phosphate buffered saline. Dilutions of the CDTb oligomeric intermediates (fractions 4, 5, 6, 7, and 8) were added to the cells at a final concentration of 7 nM, 1.4 nM, 280 pM, or 56 pM. CDTa was added to a final concentration of 10 nM for all conditions. Cellular nuclei were imaged once at the onset of the experiment in the blue channel (an excitation of 381-400 nm and emission of 414-450 nm) and cell rounding was monitored in bright field, collecting images every 20 minutes over the course of fifteen hours. All cellular samples were maintained at 5% CO_2_ at 37 ℃ over the duration of the experiment.

The total number of cells in each field was estimated as the number of nuclei in each image. Each initial image collected in the blue channel was segmented using the Weka Segmentation program (v 3.3.4) in FIJI (v1.54f) (29,30). The segmented images were then binarized and the number of particles enumerated using the “Analyze Particles” function in FIJI (30). A similar pipeline consisting of image segmentation, binarization, and particle analysis was used to enumerate round cells in all bright field images that were collected over the duration of the experiment (29,30).

### Cryo-EM Sample Preparation, Map Reconstruction, Model Building, and Refinement

To prepare samples for Cryo-EM analysis, CDTb_45-876_ was purified as described above and diluted to a final concentration of 1 µM. The sample was mixed with 1 µM CDTa_50-463_ and flash frozen in liquid ethane using a Vitrobot Mark IV. All high-resolution data were collected on a Titan Krios transmission electron microscope operating at 300 keV and equipped with a Bioquantum energy filter set to a slit width of 20 eV. Individual movies were collected with a total dose of 50 e^-^/Å^2^. The raw movie files were aligned using Patch Motion Correction in CryoSPARC (v4.5.1) (31). The defocus of the compressed images were estimated using Patch CTF Estimation in CryoSPARC and a small subset of particles were picked by hand to generate references to pick the entire dataset (31). The oligomeric intermediate structures were sorted through iterative rounds of two-dimensional and ab initio classification, forcing each particle into only one representative class. All particle sets were reconstructed and refined using non-uniform refinement in CryoSPARC (31). Particle sets resulting in maps with anisotropic resolution were truncated using the “Rebalance Orientation” function in CryoSPARC before final refinement. Molecular models were constructed from these maps in Coot (v0.9.8.1) and refined in Phenix (v20.1.1) (32,33). The quality of the maps was estimated using MolProbity (34).

### CDTb Oligomerization Assays

Purified CDTb_45-876_ was diluted to a final concentration of 100 nM and incubated at 37 °C in an attempt to induce oligomerization. Samples were taken every 10 minutes and quenched with Laemmli buffer. All samples were analyzed by SDS-PAGE stained with silver stain. To induce oligomerization, methanol (>99.8% purity, Sigma-Aldrich 179337), ethanol (100%, Decon 2701), isopropanol (>99.9% purity, Fisher Scientific A451), dimethyl sulfoxide (>99.9% purity, Sigma-Aldrich 276855), acetone (>99.5% purity, Sigma-Aldrich 179124), and polyethylene glycol 8,000 (Fisher Scientific, BP-233-100) were incubated with CDTb at concentrations of 0.08, 0.16, 0.31, 0.63, 1.25, and 2.50 M. The mixtures were incubated at 37 °C for 60 minutes and quenched with Laemmli buffer before analysis by SDS-PAGE and silver stain. The induction of oligomerization was quantified using densitometric analysis in FIJI and defined as the ratio of oligomer formed in each reaction divided by the amount of oligomer formed in the presence of water (30).

### Induction of CDTb Oligomerization via the Inclusion of CDTa

The monomeric form of CDTb was isolated via size exclusion chromatography as described above. To monitor the formation of oligomeric particles, CDTb monomer from fraction 11 was incubated at 37 °C for one hour and the sample analyzed by size exclusion chromatography. To evaluate the effect of CDTa inclusion on CDTb oligomerization, CDTa was added at a final concentration of 1 µM to a solution containing CDTb monomer from fraction 11 and incubated for one hour at 37 °C before further analysis by size exclusion chromatography. CDTa at 1 µM was incubated at 37 °C for one hour in tandem to be used as a negative control.

## Data Availability

The structural data reported here has been made available through the Electron Microscopy Data Bank (EMDB-48170, EMDB-48171, EMDB-48172, EMDB-48173, EMDB-48174, EMDB-48175, EMDB-48176, and EMDB-48177, EMDB-48178) and Protein Data Bank (PDB 9MDI, PDB 9MDJ, PDB 9MDL, PDB 9MDN, PDB 9MDP, and PDB 9MDR).

## Acknowledgements

All Cryo-EM data were generated at the Stanford-SLAC Cryo-EM Center (S^2^C^2^). This work is supported by the National Institute of Allergy and Infectious Disease (R00AI154672, MJS), the University of Minnesota Medical School, and the University of Minnesota Research Foundation. Images generated from this work were prepared using UCSF ChimeraX which was developed by the Resource for Biocomputing, Visualization, and Informatics at the University of California, San Francisco, with support from National Institutes of Health R01-GM129325.

## SUPPLEMENTAL INFORMATION

**Supporting Fig 1.**
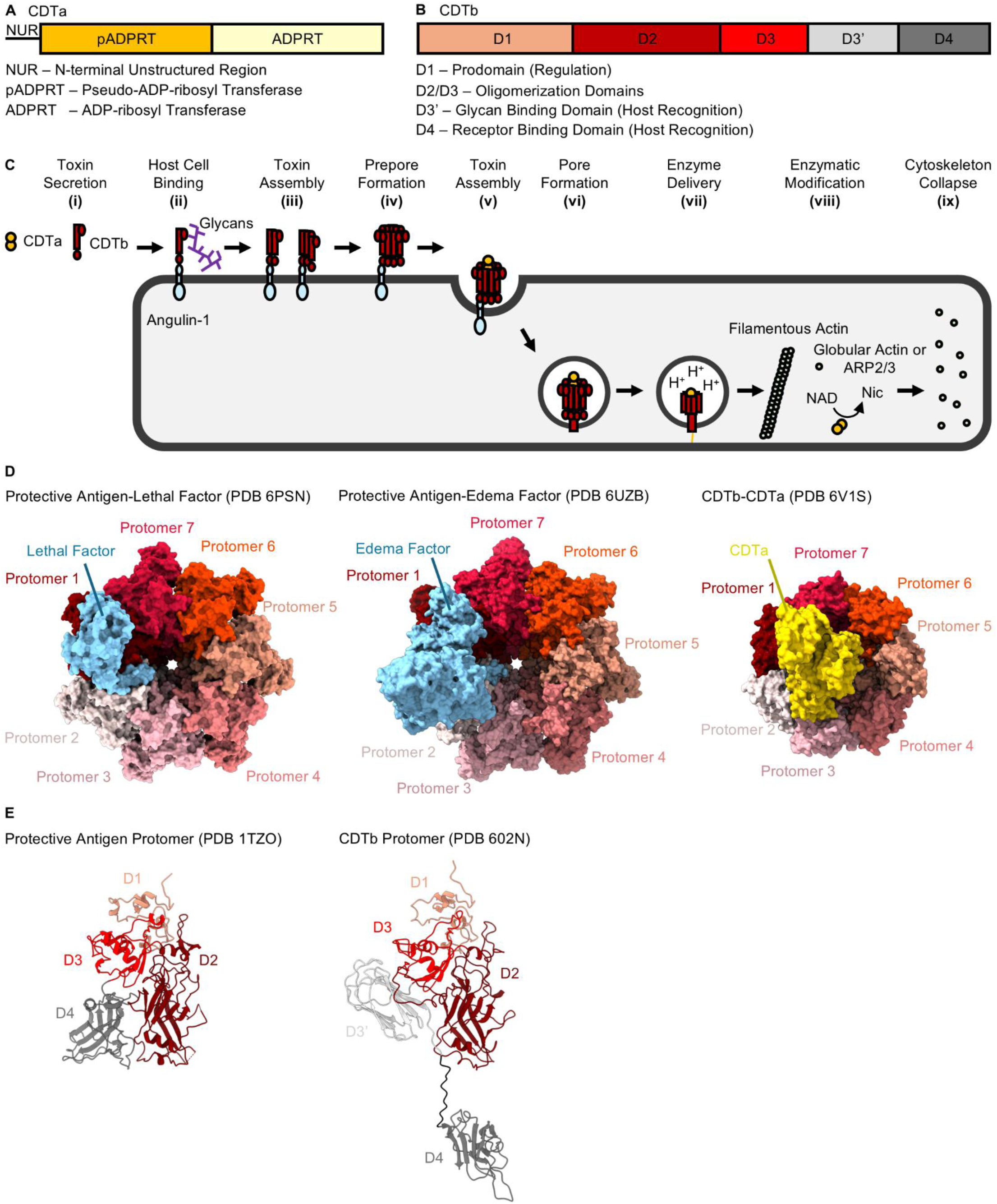
The Structure and Proposed Mechanism of CDT Intoxication. The previously defined domain organization of CDTa and CDTb are depicted in panels (**A**) and (**B**), respectively. (**C**) The accepted model of CDT intoxication begins with toxin secretion at the site of infection (i). Once secreted, CDTb localizes to host cell through a receptor known as Angulin-1. CDTb has also been shown to interact with glycans though it is not clear what role this interaction plays during intoxication (ii). CDTb is then proteolyzed and oligomerizes (iii) to form a structure referred to as the ‘prepore’ (iv). The prepore binds a single copy of CDTa and enters cells via endocytosis (v). Within the endosome CDTb undergoes a structural rearrangement leading to the formation of a membrane-spanning channel or ‘pore’ (vi). In response to the environment of the maturing endosome, CDTa passes through the CDTb pore and into the host cell cytoplasm (vii). Inside the cell, CDTa modifies globular actin and the actin related protein 2/3 (Arp2/3, viii) leading to cytoskeletal collapse and a cell rounding phenotype (ix).

**Supporting Fig 2.**
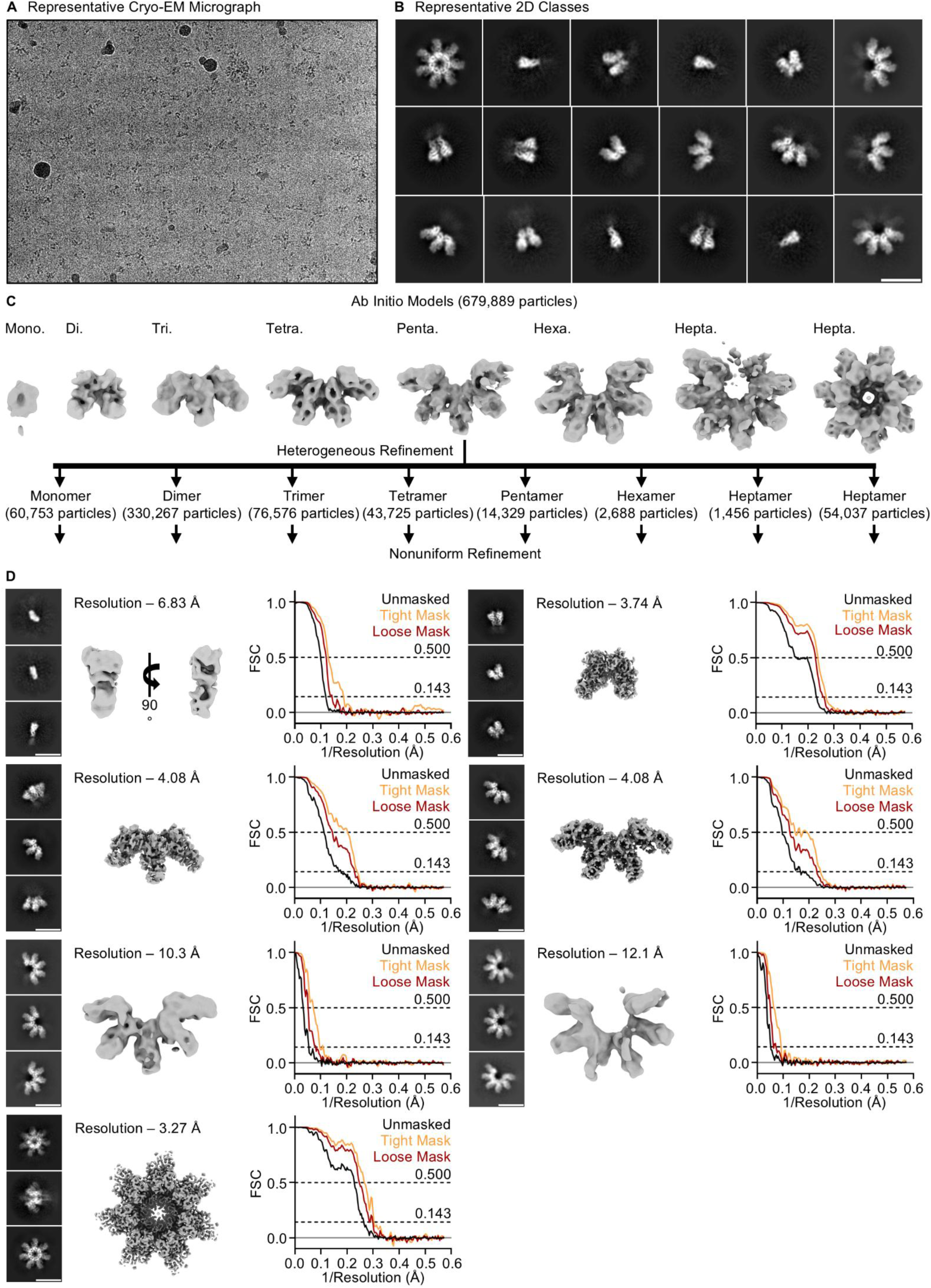
Cryo-EM Classification of CDTb Oligomeric Intermediates. (**A**) A representative micrograph collected of the sample containing CDTb oligomeric intermediates. (**B**) Two-dimensional classification was used to sort particles into distinct oligomeric states. These particles were then used to generate eight ab initio models (**C**). The ab initio models were refined to generate high resolution maps to be used as references for further refinement. All maps that were generated are shown below with representative two-dimensional classes illustrated on the left. Fourier shell correlation plots were used to estimate the global resolution of each map using a 0.143 cutoff as shown to the right of each map.

**Supporting Fig 3.**
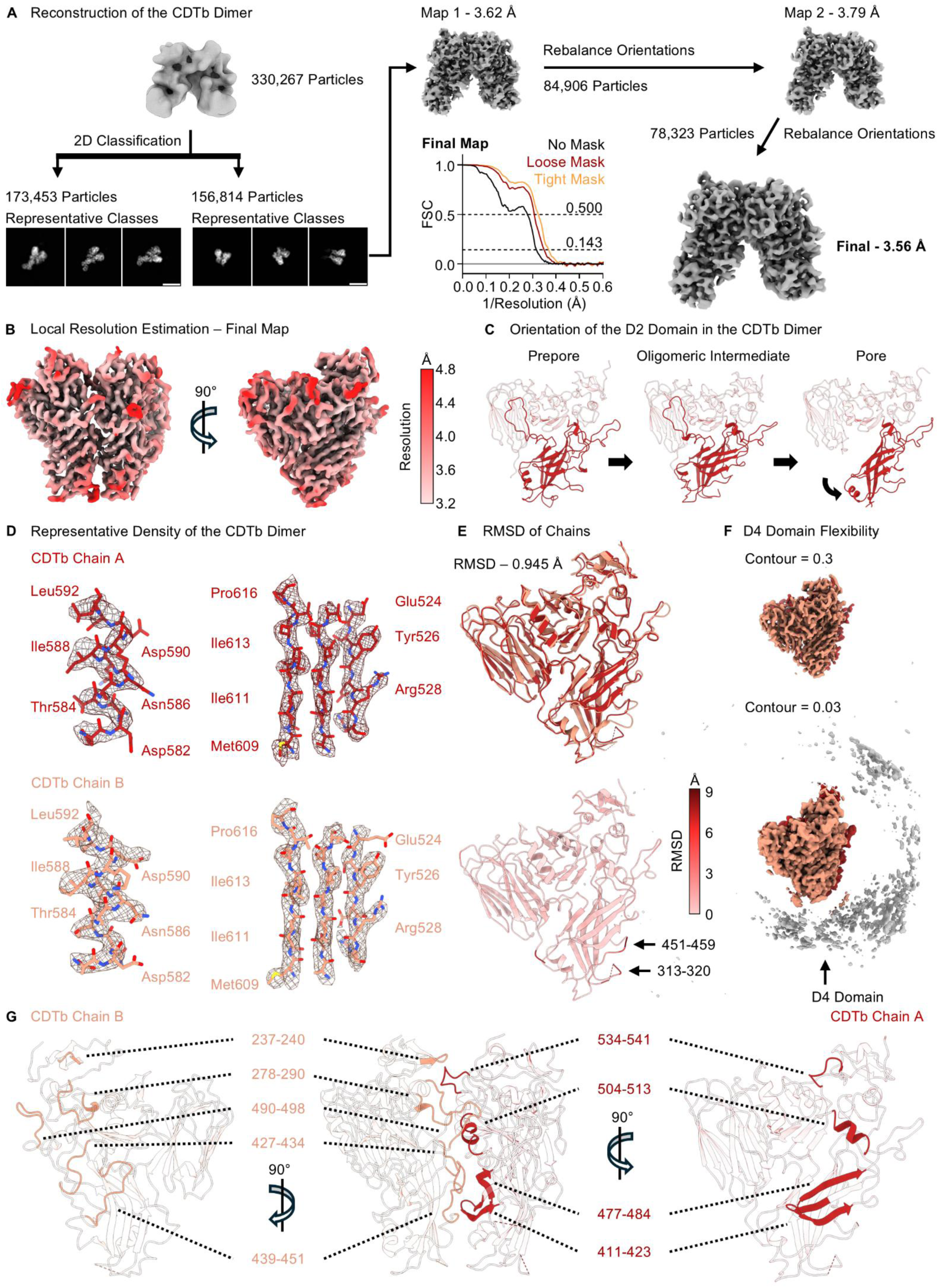
Reconstruction and Structure of the CDTb Dimeric Assembly Intermediate. (**A**) Particles corresponding to the CDTb dimer were subjected to a final round of two-dimensional classification to separate structurally distinct particles. This dataset was then used to reconstruct a map of the CDTb dimeric assembly intermediate in three-dimensional space to a global resolution of 3.62 Å. Due to issues arising from preferred orientation, the resulting map suffered from anisotropic resolution and was subjected to iterative jobs to rebalance particle orientations within the dataset. The final map was reconstructed to a resolution of 3.56 Å as determined by Fourier shell correlation using a 0.143 cutoff. (**B**) The local resolution of the CDTb dimer is displayed on the final map. (**C**) The CDTb dimeric assembly intermediate is observed in a prepore-like configuration wherein the D2 domain adopts an outward facing conformation as opposed to the inward facing conformation reported in the structure of the CDTb pore. (**D**) Representative density of Chains A and B in the CDTb dimeric intermediate. (**E**) An overlay of Chains A and B is shown at the top with the backbone RMSD of all residues observed in both structures indicated at the bottom. (**F**) A low contour map of the CDTb dimer illustrating a lack of density corresponding to the CDTb D4 receptor binding domain (top). A high contour rendering of the same map indicates the presence of density that can be attributed to the D4 domain (bottom). (**G**) Residues involved in the interface that facilitates CDTb dimerization. Chain A is shown in scarlet and Chain B in salmon.

**Supporting Fig 4.**
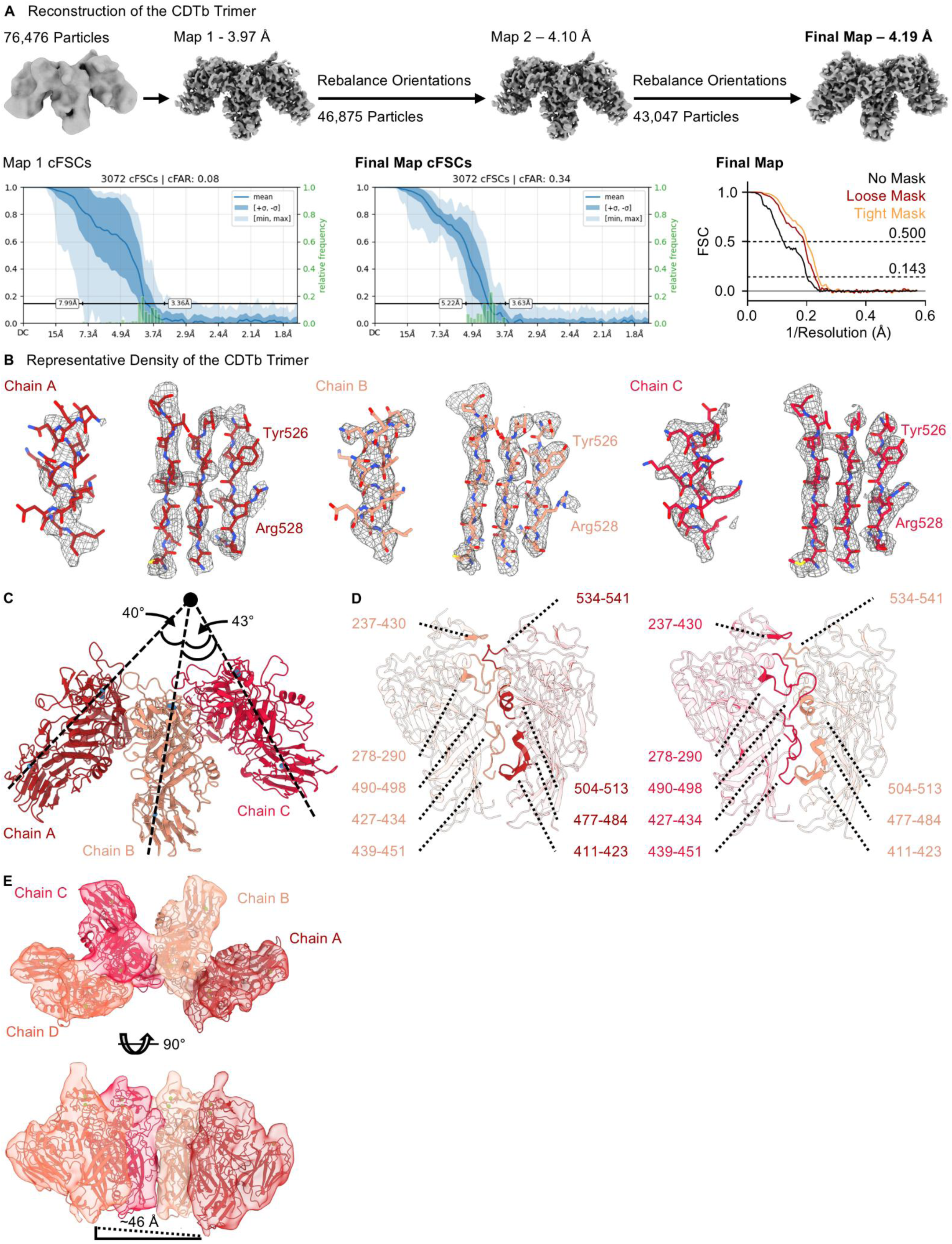
Reconstruction and Analysis of the CDTb Trimeric and Tetrameric Assembly Intermediates. (**A**) An initial map of the CDTb trimeric assembly intermediate was reconstructed to 3.97 Å resolution. Particles were rebalanced iteratively to limit the effect of preferred orientation for this sample resulting in a final map of 4.19 Å resolution. (**B**) Representative density of the CDTb trimeric intermediate with the constructed model fit into the density. Chain A is depicted in scarlet, Chain B in salmon, and Chain C in red. (**C**) The relative orientations of Chains A and B and Chains B and C with respect to the central symmetry axis of the symmetric heptamer. (**D**) Residues facilitating interactions between Chains A and B (left) and Chains B and C (right). (**E**) The low-resolution map that was reconstructed of the CDTb tetrameric assembly intermediate. Chain A is shown in scarlet, Chain B in salmon, Chain C in red, and Chain D in peach.

**Supporting Fig 5.**
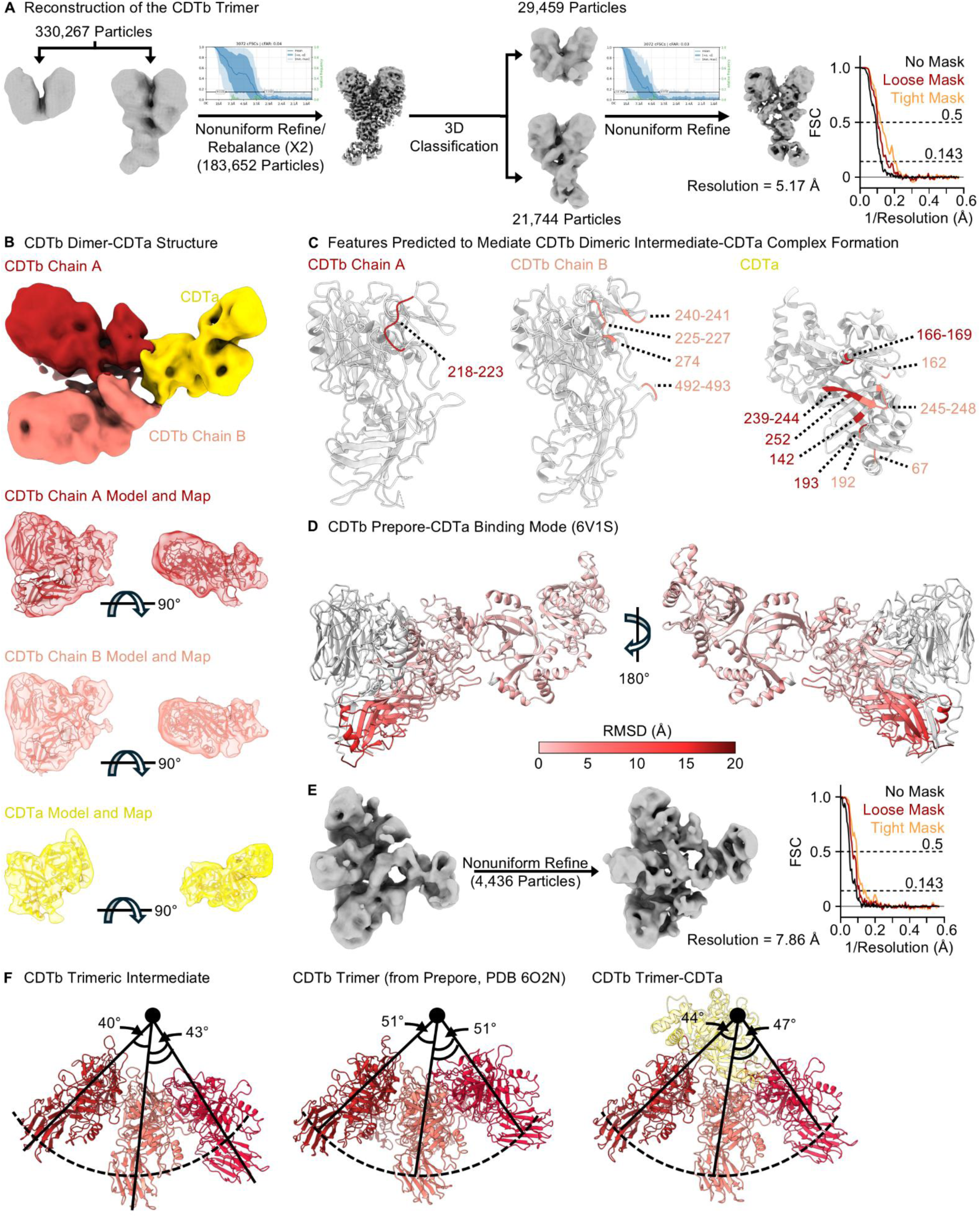
CDTa Bound Oligomeric Intermediates. (**A**) Reconstruction of the CDTb dimeric intermediate bound to CDTa resulted in a map with a global resolution of 5.17 Å. Conical FSCs (inset) were used to define the resolution of the map in three dimensions and limit the influence of preferred particle orientation in the final map. (**B**) Density of the CDTb dimeric intermediate bound to CDTa colored to illustrate the location of CDTb Chain A (scarlet), CDTb Chain B (salmon), and CDTa (gold) is shown at the top. The fit of the generated model is shown for each chain below. (**C**) Residues predicted to be at the site of interaction between the CDTb dimer and CDTa. Residues interfacing with CDTb Chain A are shown in scarlet and residues interfacing with Chain B are shown in salmon. (**D**) A plot depicting the RMSD between the CDTb dimeric intermediate bound to CDTa and CDTa bound to the CDTb symmetric heptamer (PDB 6V1S). (**E**) Reconstruction of the CDTb trimeric intermediate in complex with CDTa led to the generation of a map resolved to 7.86 Å. (**F**) The relative orientations of Chains A (scarlet), B (salmon), and C (red) are illustrated for the apo CDTb trimeric intermediate (left), the trimer structure extracted from the CDTb symmetric heptamer (middle), and the CDTb trimeric assembly intermediate bound to CDTa (right). The relative orientations of Chains A, B, and C in the CDTb trimeric assembly intermediate more closely resemble that of the CDTb symmetric heptamer.

**Supporting Fig 6.**
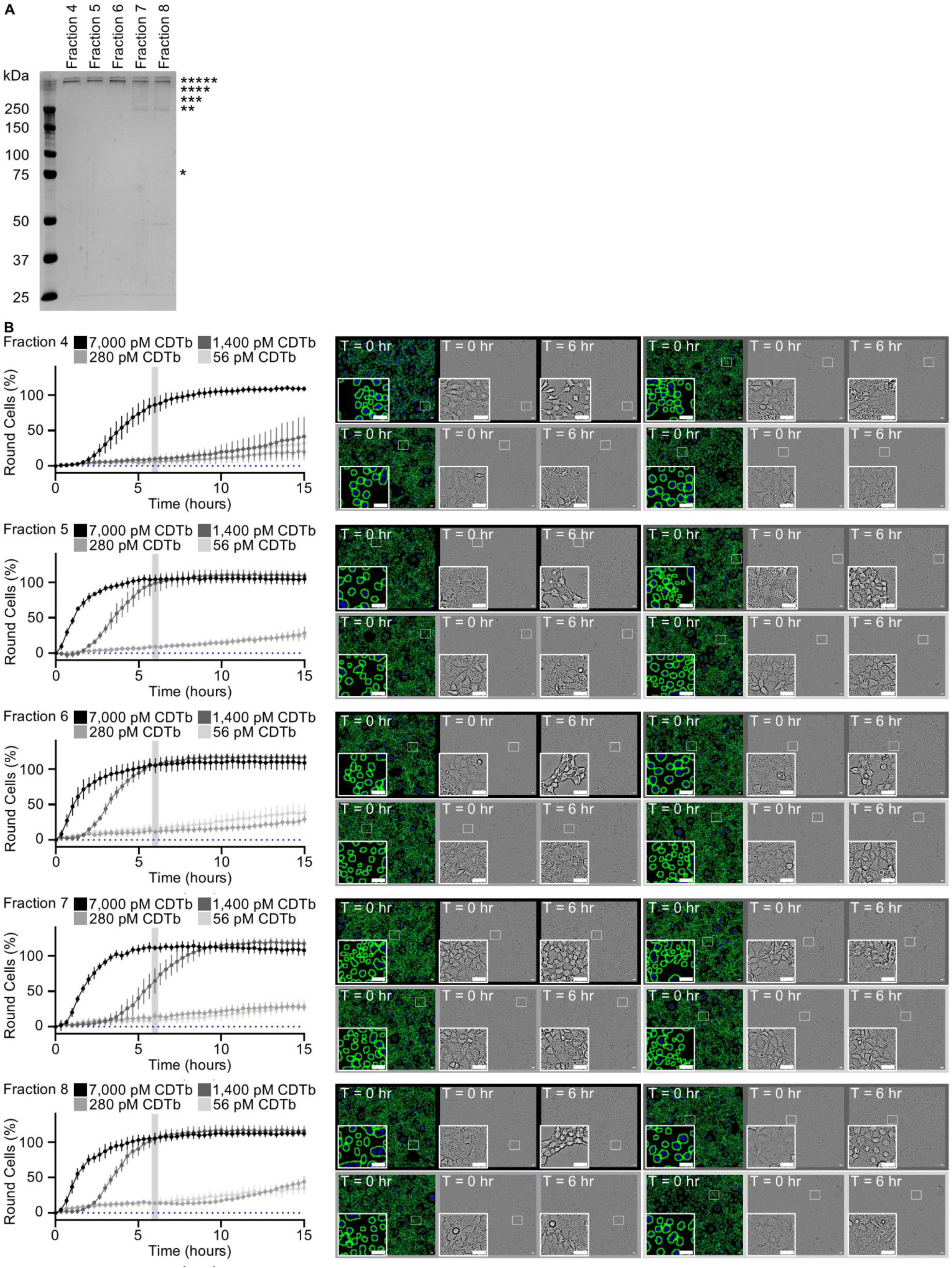
Cellular Intoxication Assays Illustrating the Effectiveness of CDTb Assembly Intermediates. (**A**) CDTb assembly intermediates were isolated via size exclusion chromatography and analyzed by SDS-PAGE. A* indicates the CDTb monomer while **, ***, ****, and ***** indicate unique molecular species with molecular weights corresponding to oligomeric assembly intermediates. (**B**) Enumeration of cellular intoxication assays for all five fractions are shown for the entire fifteen-hour assay (left). Error bars represent standard deviation. Representative images for each concentration assayed are shown on the right with the panels color-coded to reflect the concentration as depicted in the graph on the left. The first image in each series depicts the enumeration of nuclei with blue indicating Hoechst staining and green indicating the computational assignment of a nucleus. The remaining two images in each sequence are bright field images that illustrate the number of round cells at the onset of the experiment (middle) and after a six-hour incubation (right).

**Supporting Fig 7.**
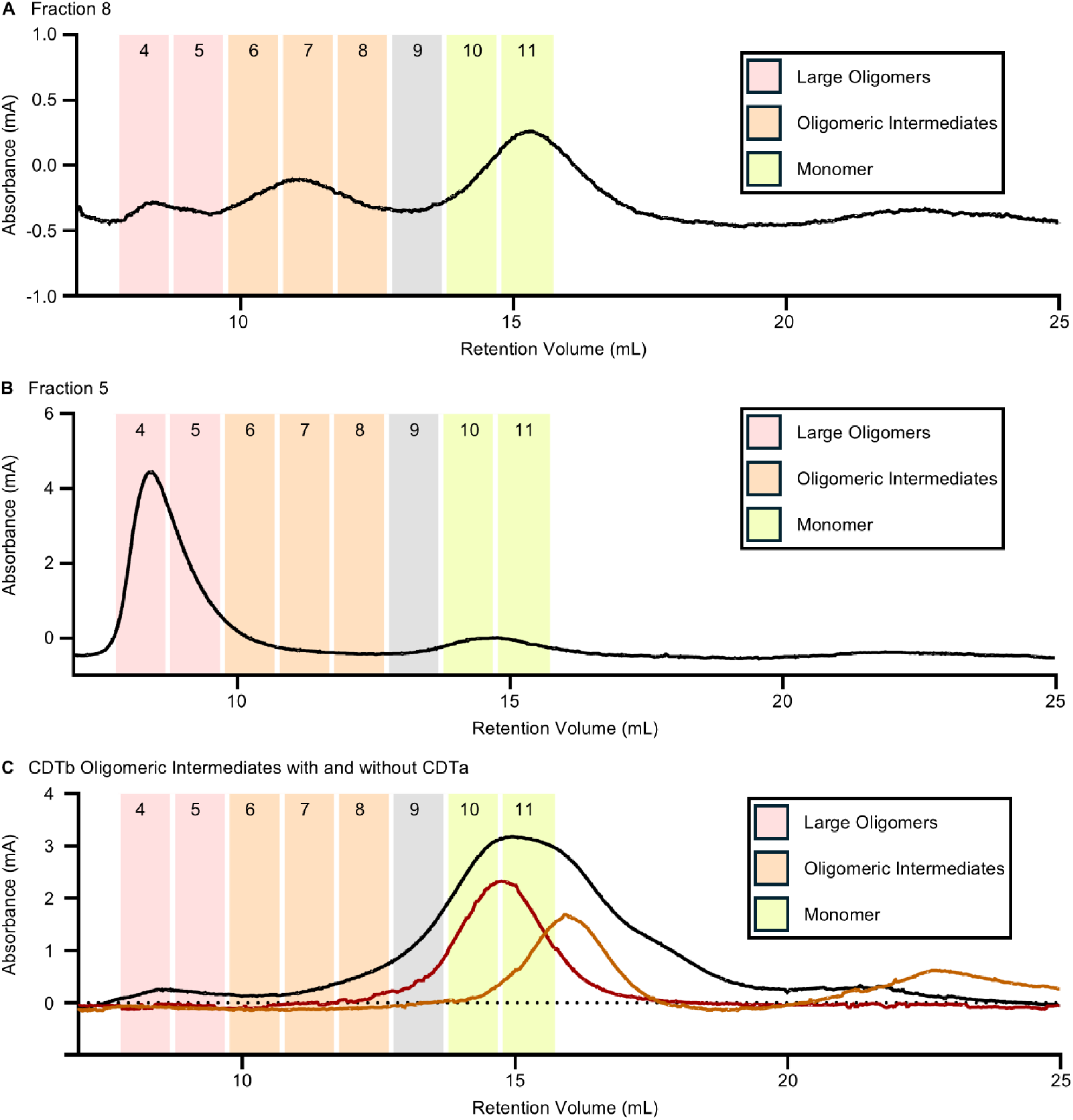
CDTb Oligomeric Intermediate Assembly is Reversible. (**A**) The content of fraction eight was assessed via size exclusion chromatography and indicates the presence of large oligomeric particles (pink), oligomeric intermediates (orange), and the CDTb monomer (yellow). (**B**) The content of fraction five was assessed via size exclusion chromatography illustrating the presence of large oligomeric particles (pink) in this fraction. No oligomeric intermediates (orange) and a relatively low abundance of the CDTb monomer (yellow) were observed in this sample. (**C**) Size exclusion chromatography analysis of CDTa, CDTb oligomeric intermediates, and CDTb oligomeric intermediates in the presence of CDTa.

**Supporting Fig 8.**
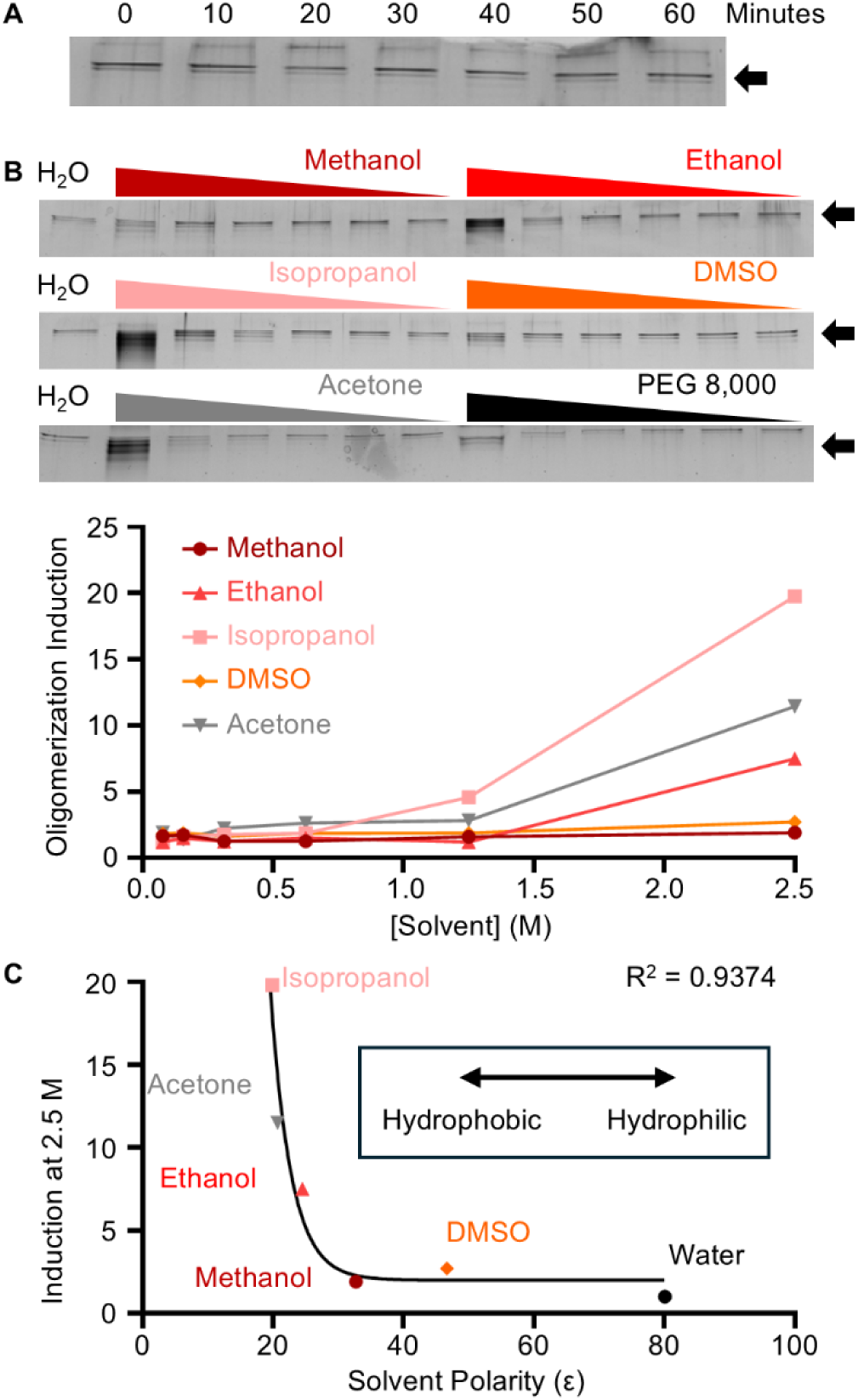
Induction of CDTb Oligomerization by Hydrophobic Solvents. (**A**) In agreement with previous studies we note the oligomerization of CDTb is slow in vitro with little oligomer formation occurring after a 60-minute incubation. (**B**) Oligomerization was induced through the addition of various solvents and polyethylene glycol 8,000 (PEG 8,000). The concentration of the oligomer was quantified at various concentrations. DMSO – dimethyl sulfoxide. (**C**) The induction of oligomerization at 2.5 M solvent plotted against the polarity of the solvent and fit to an exponential decay function. DMSO – dimethyl sulfoxide.

**Supporting Table 1.**
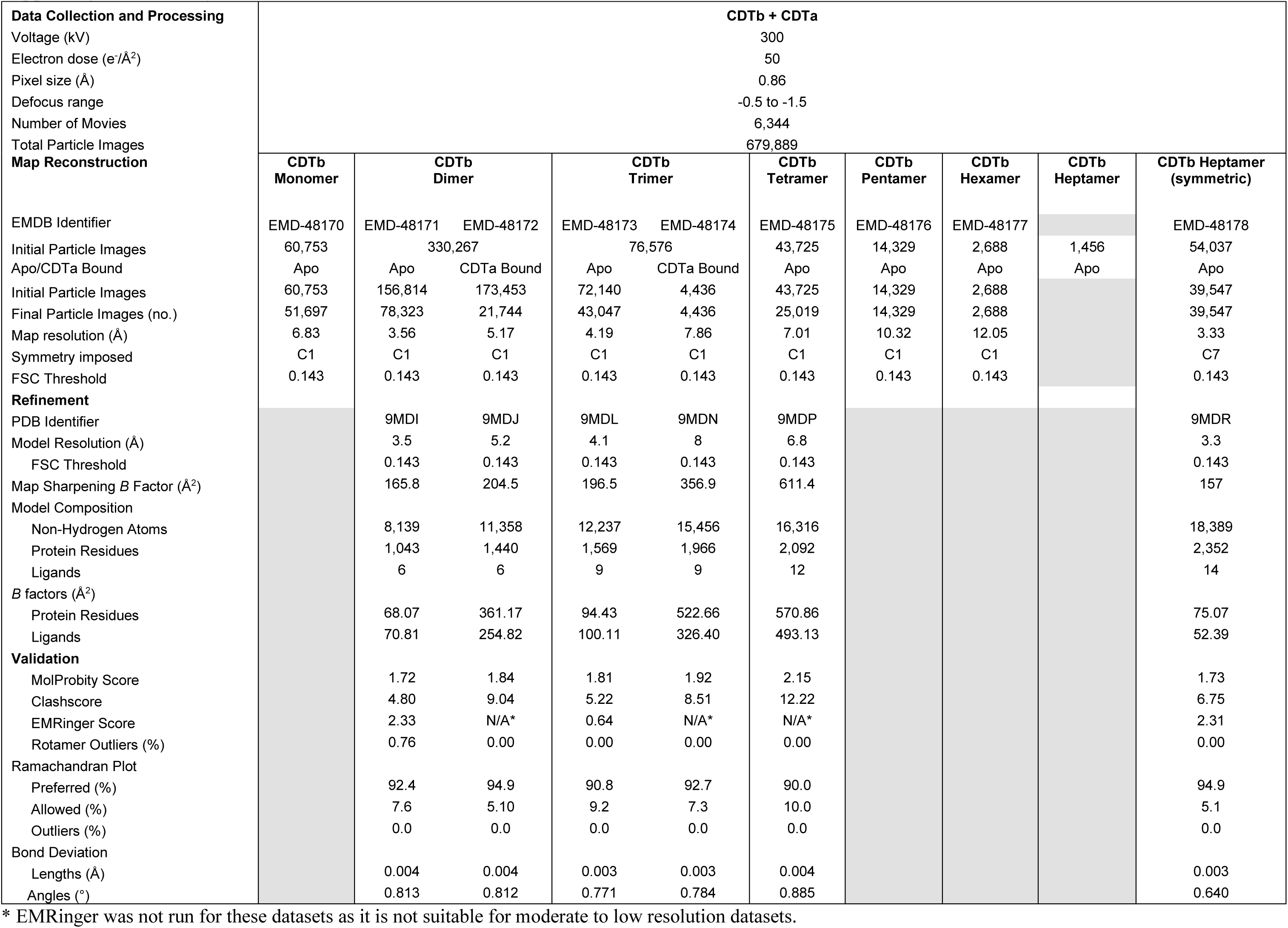
Cryo-EM Data Collection and Refinement Statistics.

